# Sediment accumulation, elevation change, and the vulnerability of tidal marshes in the Delaware Estuary and Barnegat Bay to accelerated sea level rise

**DOI:** 10.1101/821827

**Authors:** LeeAnn Haaf, Elizabeth Burke Watson, Tracy Elsey-Quirk, Kirk Raper, Angela Padeletti, Martha Maxwell-Doyle, Danielle Kreeger, David Velinsky

## Abstract

Tidal marshes protect coastal communities from the effects of sea level rise and storms, yet they are vulnerable to prolonged inundation and submergence. Uncertainty regarding their vulnerability to sea level rise motivated the establishment of a monitoring network in the Delaware Estuary and Barnegat Bay. Using data collected through these efforts, we determined whether rates of tidal marsh sediment accumulation and elevation change exceeded local sea level rise and how these dynamics varied along geographic and environmental gradients. Marker horizons, surface elevation tables, elevation surveys, water level data, and water column suspended sediment concentrations were used to evaluate sea level rise vulnerability. Of 32 study sites, 75% had elevation change that did not keep pace with long-term rising sea levels (1969–2018) and 94% did not keep pace with recent sea level rise (2000–2018). Mean high water rose most rapidly in the freshwater tidal portion of the Delaware Estuary with rates nearing 1 cm yr^-1^ from 2000–2018. We noted that greater sediment accumulation rates occurred in marshes with large tidal ranges, low elevations, and high water column suspended sediment concentrations. We found correlations between rates of shallow subsidence, increasing salinity, and decreasing tidal range. Marsh elevation and water level surveys revealed significant variability in elevation capital and summer flooding patterns (12–67% inundation). However, rapid increases in mean high water over the past 19 years suggests that all marsh platforms currently sit at or below mean high water. Overall, these data suggest that tidal marshes in the Delaware Estuary and Barnegat Bay are vulnerable to submergence by current rates of sea-level rise. While we observed variability in marsh elevation capital, the absence of strong correlations between elevation trends and environmental parameters makes it difficult to identify clear patterns of sea level rise vulnerability among wetlands.

## Introduction

Tidal marshes can moderate some of the impacts of climate change including storm surge and wave attenuation, nutrient uptake and removal through denitrification, and mitigation of greenhouse gas emissions through carbon sequestration [1–3]. An analysis of damage caused by Hurricane Sandy in the U. S. Mid-Atlantic suggested that intact tidal marshes reduced flood damages by more than US $625 million and lower annual flood risks around Barnegat Bay, NJ by up to 70% [4]. Ensuring the existence of tidal marshes in coastal areas bordering high-value infrastructure over the coming decades is an important part of protecting coastal communities from increased flood risk due to sea level rise (SLR) and extreme weather events like hurricanes [4–6].

Despite being assets for coastal community protection, tidal marshes are vulnerable to the impacts of SLR, especially when additional anthropogenic stressors reduce their elevation or accretionary capacity, such as declining sediment inputs and altered hydrology [7–10]. Accumulation of plant material in the soil and sediment accumulation interact to build elevation through dynamic feedbacks with sea level [11–16]. Declines in sediment availability caused by channel dredging, upstream damming, and changes in agricultural practices, coupled with hydrological changes, such as mosquito ditching, attenuate tidal marsh responses to sea level changes by reducing sedimentation rates and increasing inundation [9, 17,18]. Further, as rates of SLR increase, it is not clear whether marshes that may already be vulnerable to inundation and low sediment supply will be able to increase elevation at rates that would prevent submergence.

SLR in the U. S. Mid-Atlantic is increasing faster than the global average due to steric sea level changes and geologic subsidence [19–23]. Long-term observations derived from local National Oceanic and Atmospheric Administration (NOAA) tidal gauges show that SLR in Delaware Estuary (ranging from 2.98 ± 0.19 to 4.63 ± 0.50 mm yr^-1^) and Barnegat Bay (4.09 ± 0.15 mm yr^-1^) are nearly twice that of the early 20th century global average (1.7 mm yr^-1^)[23]. Projections of local SLR in Delaware suggest that rates will likely exceed 10 mm y^-1^ by 2100 for intermediate or high emission scenarios [24]. This projected rate approaches a critical threshold for tidal marsh elevation feedbacks, and suggests marsh drowning will occur [16]. Additionally, subsidence driven by local groundwater withdrawal [25] or historical land manipulations [26], such as diking, accelerates SLR locally and further expedites marsh drowning. In fact, tidal marsh loss due to submergence across the U. S. Northeast and Mid-Atlantic is already widespread [27–29].

Extensive disturbances in Barnegat Bay and below average elevations relative to tides in the Delaware Bay suggests that tidal marshes in both estuaries have a high degree of vulnerability to accelerating rates of SLR [18, 30–36]. Tidal marsh losses and increased interior flooding have been documented in both estuaries [7, 26–27, 30–36], but no analyses of elevation dynamics across these areas have been completed to date. Further, elevation change data from other locations in the Mid-Atlantic and Northeast have shown that SLR frequently exceeds marsh accumulation [37–42].

The importance of maintaining Mid-Atlantic tidal marshes for community protection, combined with ongoing evidence of marsh drowning, motivated the establishment of a monitoring network. Two National Estuary Programs in collaboration with the Academy of Natural Sciences of Drexel University established this network to determine how tidal marshes in the Delaware Estuary and Barnegat Bay are responding to accelerated SLR (www.macwa.org). A variety of data were collected as part of these monitoring efforts, but this particular study focuses on surface elevation change and surface accretion rates. Our objectives were to: (1) compare rates of marsh sediment accumulation and vertical elevation change to long-term and contemporary rates of SLR, as well as contemporary rates of rise in mean high water for the Delaware Estuary and Barnegat Bay; (2) to determine how elevation capital, salinity, tidal range, and water column suspended sediments influence elevation dynamics among geographically distinct tidal marshes.

## Materials and methods

### Study Sites

The Delaware Estuary, located in New Jersey, Delaware, and Pennsylvania, contains over 66,000 ha of tidal marsh [35]. It has a length of approximately 215 km from the fall line at Trenton, New Jersey to its confluence with the Atlantic Ocean. Upstream of Wilmington, Delaware, the estuary is freshwater (<0.3‰) riverine and has a tidal range between 1.6 and 2.3 m. Downstream of Wilmington, salinities gradually transition to polyhaline (20–25‰) with tidal ranges from 1.2 to 1.7 m. Freshwater, brackish, and saline tidal marshes exist along this estuarine gradient. *Zizania aquatica, Peltandra virginica, Polygonum punctatum*, and *Nuphar advena* are among many herbaceous perennial and annual species that dominate tidal freshwater marshes in the Delaware River. Halophytic grasses, such as *Spartina alterniflora* (syn. *Sporobolus alterniflorus*), *Spartina patens* (syn. *Sporobolus pumilus*) and *Distichlis spicata*, dominate the brackish and saline marshes of the Delaware Bay. Because of heavy development in the Trenton-Philadelphia-Wilmington industrial corridor, only 16.7% of tidal freshwater wetlands remain in Pennsylvania from the pre-colonization extent (500 of ∼3,000 ha) [35, 43] and at least six fresh intertidal plant species are extirpated from the tidal Delaware River [44]. Satellite imagery analyses suggest that marsh loss in the Delaware Estuary was ∼80 ha y^-1^ from 1996–2010 [35]. Although wetland loss to development has declined, major threats remain. Marsh loss in the Delaware Estuary are occurring due to open water conversion linked to historical manipulations (*e*.*g*. diking, ditching), pervasive shoreline erosion, and anthropogenic limitations to sediment supplies (*e*.*g*. dredge spoil disposal, changes in agricultural practices, damming) [9, 26, 35, 44].

Barnegat Bay is a 520-km^2^ shallow lagoon bordered to the east by a 65 km long barrier island separating the bay from the Atlantic Ocean. The 9,200 ha of salt marsh [36] fringing the Barnegat Bay are polyhaline (>18‰) and dominated by *S. alterniflora, S. patens*, and *D. spicata*. Tidal ranges are approximately <0.2–0.7 m depending on bathymetry and distance from the Bay’s two inlets [45]. While northern Barnegat Bay is largely urbanized with extensive shoreline hardening (e.g. bulkheads), the southern region is less developed and contains more protected lands (i.e., the Edwin B. Forsythe National Wildlife Refuge holds ∼14,800 ha) [46]. Estimated rates of salt marsh loss in the Barnegat Bay were ∼6 ha y^-1^ from 1996–2010 [36]. Tidal marshes manipulation for mosquito population control have been extensive, with many marshes grid ditched and over 7,000 mosquito management ponds excavated [47]. Open marsh water management (OMWM), a mosquito management tactic, increases pond habitat to lure insectivorous fish to the high marsh, but the process also causes vegetation losses and affects elevation, as excavated peat is side-casted onto the marsh [18, 47].

### Monitoring protocols

We established eleven monitoring sites in tidal marshes of the Delaware Estuary and Barnegat Bay, which varied in tidal range, salinity, and dominant vegetation type (Fig 1; Table 1). For each site, three deep rod surface elevation tables (SET) were installed [48–49] between 2010 and 2014 to measure marsh elevation change. Three feldspar marker horizon (MH) plots encircle each SET to measure short-term vertical sediment accumulation, or accretion [50]. Combined, the SET-MH technique distinguishes elevation change by surface and subsurface processes, and can provide estimates of resilience to SLR [42, 49, 51]. Here, we calculated shallow subsidence by subtracting elevation change from accretion rates, so that negative values represent declining subsurface elevations [51]. We read SET-MHs twice a year over the course of 5–7 years, depending on installation dates.

**Table 1.** Site descriptions for the Delaware River, Delaware Bay, and the Barnegat Bay.

| Region | Site | Install Year | Location | Latitude, Longitude | Tidal Range (m) | Salinity (‰) | Dominant Vegetation |
| --- | --- | --- | --- | --- | --- | --- | --- |
| Delaware River | Crosswicks Creek | 2011 | Bordentown, NJ | 40° 9.76'N, 74° 42.51'W | 2.4 | 0.10 | *Z. aquatica*, *P. virginica*, *P.* |
|  |  |  |  |  |  |  | *punctatum*, and *N. advena* |
|  | Tinicum | 2010 | Philadelphia, PA | 39° 53.05'N, 75° 16.49'W | 1.7 | 0.18 | *Z. aquatica*, *P. virginica*, *P. punctatum*, and *N. advena* |
|  | Christina River | 2010 | Wilmington, DE | 39° 43.29'N, 75° 34.07'W | 1.7 | 0.40 | *Typha angustifolia*, *P. virginica*, *P. punctatum*, and *N. advena* |
| Delaware Bay | Dividing Creek | 2012 | Dividing Creek, NJ | 39° 14.14'N, 75° 6.76'W | 1.7 | 17 | *S. alterniflora*, *S. patens*, and *D. spicata* |
|  | Maurice River | 2010 | Bivalve, NJ | 39° 15.95'N, 74° 59.72'W | 1.7 | 11 | *S. alterniflora*, *S. patens*, and *D. spicata* |
|  | Dennis Creek | 2011 | South Dennis, NJ | 39° 10.58'N, 74° 51.74'W | 1.6 | 16 | *S. alterniflora*, *S. patens*, and *D. spicata* |
|  | Broadkill River | 2014 | Lewes, DE | 39° 47.13'N, 75° 10.75'W | 1.1 | 27 | *S. alterniflora*, *S. patens*, and *D. spicata* |
| Barnegat Bay | Island Beach | 2011 | Seaside Park, NJ | 39° 47.96'N, 74° 6.10'W | 0.2 | 27 | *S. alterniflora* |
|  | Dinner Point Creek | 2010 | West Creek, NJ | 39° 37.43'N, 74° 16.20'W | 0.6 | 26 | *S. alterniflora* |
|  | Horse Point | 2012 | West Creek, NJ | 39° 37.59'N, 74° 15.43'W | 0.6 | 26 | *S. alterniflora* |
|  | Reedy Creek | 2010 | Brick, NJ | 40° 1.74'N, 74° 5.07'W | 0.2 | 20 | *S. alterniflora*, *S. patens*, and *D. spicata* |

**Fig 1.**
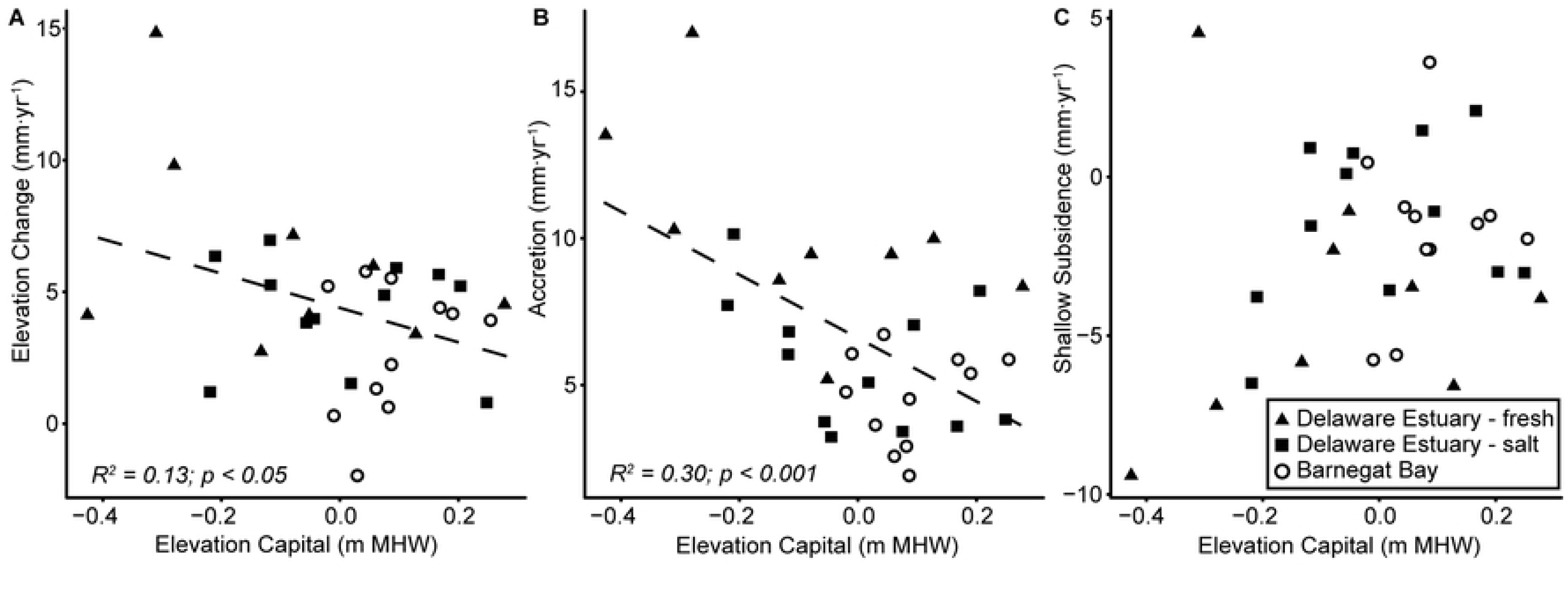
Monitoring sites in the tidal marshes of the Delaware Estuary and Barnegat Bay.

We conducted topographic surveys of tidal marsh elevations in each location from 2014 to 2018 using real-time kinematic GPS receivers (a Leica Viva GS14 GNSS Receiver and Viva CS15 field controller, or a Trimble R10 GNSS receiver and TSC2 data controller). Data collection followed National Geodetic Survey guidelines for the RT3 accuracy class (0.04–0.06 m horizontal; 0.04–0.08 m vertical precision): baselines <20 km, collection at 1 s intervals for 15s, with a steady fixed height rover pole without use of a bipod [52].

We measured a suite of physicochemical parameters along five water quality stations 2–3 times per year, in spring, fall, and/or summer. For each station, salinity was measured with a YSI (Professional Pro+, Xylem, Yellow Springs, OH) and samples were collected for further laboratory analyses, including total suspended sediment and dissolved nutrient concentrations (not reported). Additionally, we used existing available data and short-term water level logger deployments to determine local tidal datums for each site (Supporting Information).

### Calculations and data analysis

Elevation capital, often expressed as marsh elevation relative to a tidal datum such as mean high water (MHW), is a useful proxy for tidal marsh vulnerability to SLR [14]. Tidal marshes with elevations near the upper limit of intertidal plant growth possess greater elevation capital and are more resilient to rising sea levels. Tidal datums relative to the National Tidal Datum Epoch (NTDE; 1983–2001) were computed for eight sites using the modified-range-ratio method on short-term water level measures derived from in-situ loggers (Supporting Information) [53]. This method is generally associated with accuracy of 2–3 cm for a 5-month period [54]. NOAA tide stations of Atlantic City, NJ, Lewes, DE, Cape May, NJ, or Philadelphia, PA served as controls. For Crosswicks Creek and the Christina River, tide ranges matched nearby NOAA tide stations (Newbold, PA and Delaware City, DE). Because these stations lacked datum conversions to the North American Vertical Datum 1988 (NAVD88), we obtained reference water levels by surveying water levels with real time kinematic surveys to permit conversion. We calculated average flooding times (%) for the median marsh elevations using 2016 or 2018 water level data. Elevation capital, expressed relative to MHW, was the difference between median survey elevation (m NAVD88) and the elevation of local MHW (m NAVD88) relative to the National Tidal Datum Epoch (1983–2001).

To estimate rates of tidal marsh elevation change and sediment accumulation across the Delaware Estuary and Barnegat Bay, we constructed regressions of elevation and accretion against time [48, 51, 55–56]. We compared rates of accretion and vertical elevation change with local sea level rise (LSLR) trends derived from the nearest NOAA tide station [57]. We also compared trends in rates of elevation gain and sediment accretion with long-term (LT) (1969– 2018) and short-term (ST) (2000–2018) trends in SLR and, as well as ST (2000–2018) rates of increase in mean high water level (ST MHWR). To be conservative, we concluded that a site was gaining elevation or accreting sediment at rates significantly greater than SLR if its rate of rise (±SE) exceeded LSR. Associations were identified between key processes that determine marsh survival (elevation capital, elevation change, accretion, and shallow subsidence) with gradients in environmental conditions (salinity, tidal range, water column dissolved organic carbon, and suspended sediment concentration) using linear regression.

Between 2011 and 2013, pond excavation associated with OMWM affected all of the SET-MHs at Dinner Point Creek in Barnegat Bay. We began monitoring in 2010-2012, prior to pond construction. In 2012, these activities casted ∼0.20 m of sediment directly onto SET 1 (Supporting Information). We then installed SET-MHs at Horse Point, an unaffected area adjacent to Dinner Point Creek. We excluded Dinner Point Creek SET 1 data from some analyses, where noted, because they were not representative of natural or ambient processes.

## Results

Marsh elevation capital relative to MHW was variable among focal tidal marshes (Fig 2). In the Delaware Estuary, marshes along the Broadkill River, the Christina River, and Dennis Creek possessed notable elevation capital, whereas Tinicum marshes sat low in the tidal frame. In Barnegat Bay, Reedy Creek and Island Beach had lower elevation capital compared to Dinner Point Creek and Horse Point. We found significant differences between long-term (i.e. NTDE) and current high tide flooding levels (Tables 2 and 3), suggesting most marshes, with the exception of Dinner Point Creek and Horse Point, sit well below the current MHW. Elevation of the marsh relative to MHW influenced accretion (*R*^2^=0.30, *p*<0.001) and elevation change (*R*^2^=0.13, *p*<0.05), such that sediment accretion and elevation change rates were greater for marshes sitting low in the tidal frame (Fig 3). However, elevation did not influence rates of shallow subsidence.

**Table 2.** Median tidal marsh elevations relative to NAVD88, local MHW relative to the NTDE, and summer-fall mean high tide levels. Values were derived from 2016 water level data, with the exception of the Broadkill River, where data are from 2018.

| Site | Median Elevations (m) |  |  | % Time Inundated |
| --- | --- | --- | --- | --- |
|  | NAVD88 | MHW NTDE | MHW 2016 |  |
| Crosswicks Creek | 1.10 | -0.18 | -0.29 | 20% |
| Tinicum | 0.44 | -0.37 | -0.53 | 39% |
| Christina River | 0.62 | 0.07 | -0.34 | 30% |
| Dividing Creek | 0.68 | -0.04 | -0.29 | 27% |
| Maurice River | 0.67 | -0.05 | -0.30 | 25% |
| Dennis Creek | 0.61 | -0.03 | -0.21 | 24% |
| Broadkill River<sup>1</sup> | 0.39 | 0.02 | -0.22 | 27%<sup>1</sup> |
| Reedy Creek | 0.13 | 0.05 | -0.21 | 67% |
| Island Beach | 0.19 | 0.13 | -0.09 | 42% |
| Horse Point | 0.40 | 0.18 | -0.01 | 12% |
| Dinner Point Creek | 0.44 | 0.25 | 0.00 | 12% |

**Table 3.** Long (1969–2018) and short-term (2000–2018) water level trends at NOAA harmonic tidal datum stations from linear regression of monthly mean tidal data.

| Tide gage | Associated monitoring site(s) | Mean sea level rise (mm yr<sup>-1</sup>) |  | Mean high water rise (mm yr<sup>-1</sup>) |
| --- | --- | --- | --- | --- |
|  |  | LT 1969–2018 | ST 2000–2018 | ST 2000–2018 |
| Philadelphia, PA | Crosswicks Creek,<br>Tinicum | 4.03 | 6.39 | 9.53 |
| Reedy Point, DE | Christina River | 4.16 | 5.60 | 7.11 |
| Lewes, DE | Broadkill River | 3.96 | 6.72 | 7.77 |
| Cape May, NJ | Dividing Creek,<br>Maurice River, Dennis<br>Creek | 4.86 | 6.81 | 8.53 |
| Atlantic City, NJ | Reedy Creek, Island<br>Beach, Horse Point,<br>Dinner Point Creek | 4.69 | 5.75 | 7.71 |

**Fig 2.**
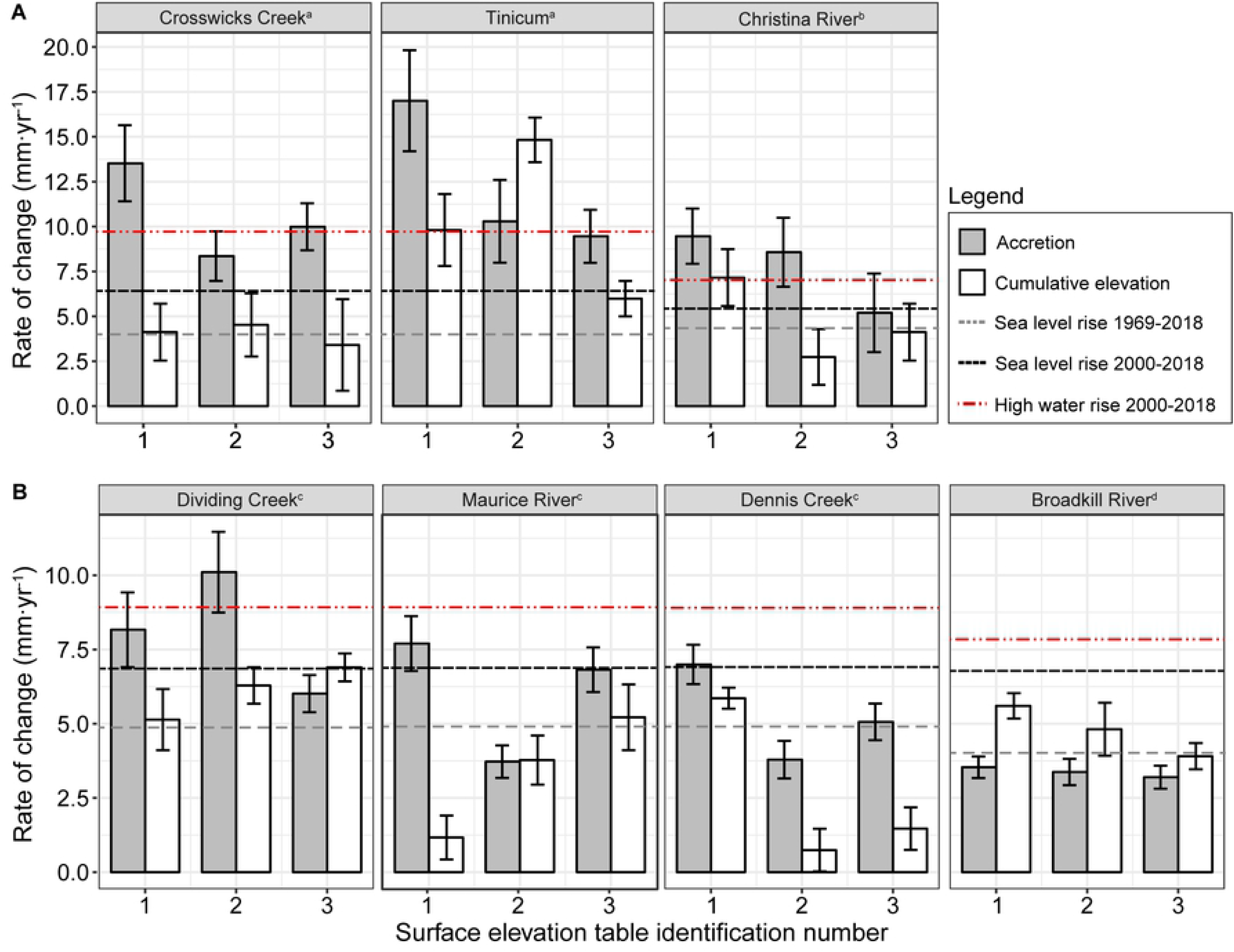
Frequency distribution of marsh elevations in Barnegat Bay and the Delaware Estuary relative to local MHW (NTDE) in meters. Vertical dashed line is 0.0 m MHW.

**Figure 3.**
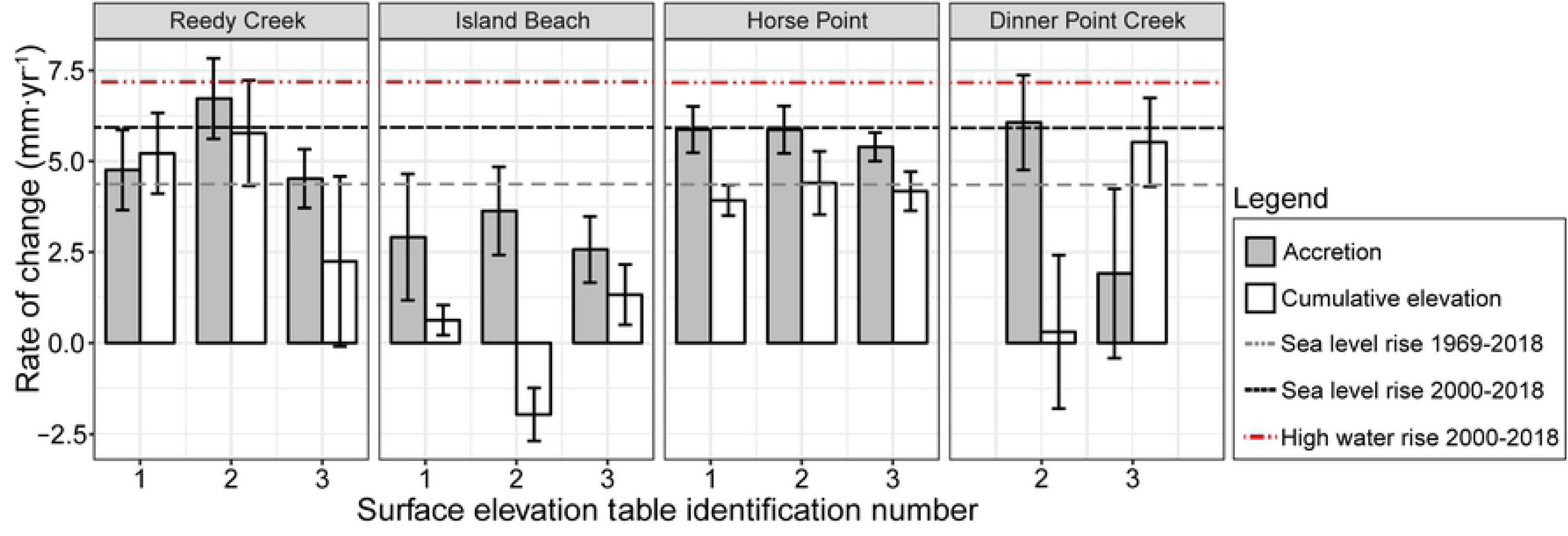
Relationship between elevation capital relative to 2016 MHW and (A) elevation change, (B) accretion, and (C) shallow subsidence. Solid triangles are freshwater tidal marshes in the Delaware River, solid squares are salt marshes in the Delaware Bay, and hollow circles are salt marshes in the Barnegat Bay. There were no significant differences among estuaries.

Rates of LSLR and MHWR varied depending on the time period under consideration, with lower rates associated with longer time windows, and higher rates recorded over the past 19 years (Table 3). Notably, LSLR was 44% greater over the past 19 years than over the last 50 years, and over the past 19 years, the rise in MHW was significantly greater (30%) than LSLR (Table 3). The most rapid rate of rise was found for tidal marshes in the freshwater tidal portion of the Delaware Estuary, where MHWR has approached 1 cm yr^-1^ (0.95 cm yr^-1^), values that exceed even the most extreme forecasts for 21^st^ century SLR rates (Najjar et al. 2000).

Comparisons of accretion and elevation change rates with LT LSLR show variability within and across estuaries and sites. Of the 32 surface elevation tables, excluding Dinner Point Creek SET 1, 75% had elevation trends that did not exceed rates of LT LSLR (1969–2018) (Supporting Information). When compared to ST LSLR and MHWR, only 6% and 4% of marsh areas had rates of elevation increase that kept pace, respectively. Accumulation deficits, or the difference between LT LSLR and elevation change, varied from −4.12 to 10.8 mm yr^-1^ in the Delaware Estuary and −6.65 to 1.08 mm yr^-1^ in Barnegat Bay (Table 4) (Supporting Information).

**Table 4.**
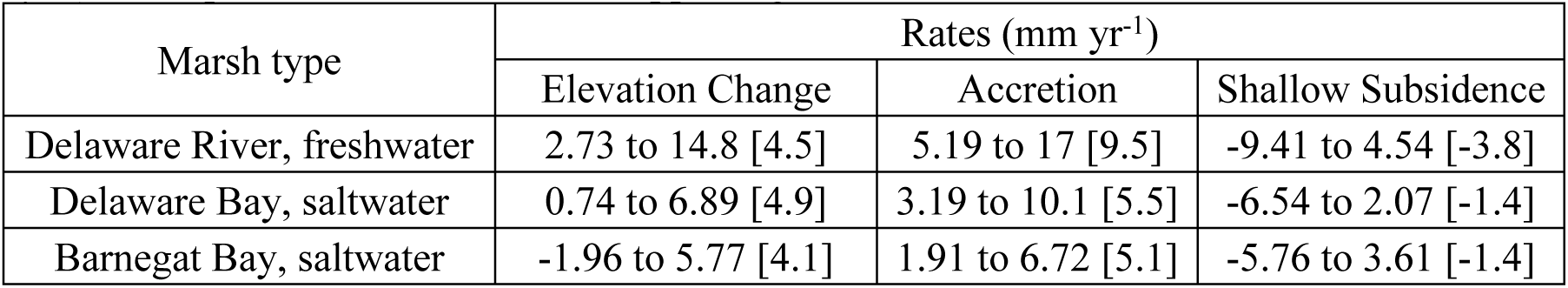
Ranges of elevation change, accretion, and shallow subsidence summarized by tidal freshwater marshes in the Delaware River, salt marshes in the Delaware Bay, and salt marshes in Barnegat Bay. Median values are in brackets. In Barnegat Bay, Dinner Point Creek SET 1 was excluded from this table, due to dispersal of sediment from mosquito management practices causing anomalously high annual mean accretion and elevation change rates (>25 mm yr^-1^). Site-specific information is in Supporting Information.

Processes of elevation change, accretion, and shallow subsidence varied between the two estuaries, as well as along the salinity gradient in the Delaware Estuary (Figs 4 and 5; Table 4). Intra-site variation of elevation change was large at some sites, with the largest range at Tinicum (8.82 mm yr^-1^). Twenty-four of the 32 (75%) SET-MHs experienced shallow subsidence (Supporting Information; excluding Dinner Point Creek SET 1). Seven (21%) SET-MHs experienced subsurface expansion and only one site (3%) had elevation changes that were loosely equivalent to accretion (Maurice SET 2). Shallow subsurface processes had a substantial role in elevation dynamics, whether it was due to subsurface expansion (presumably linked to belowground production) or consolidation.

**Fig 4.**
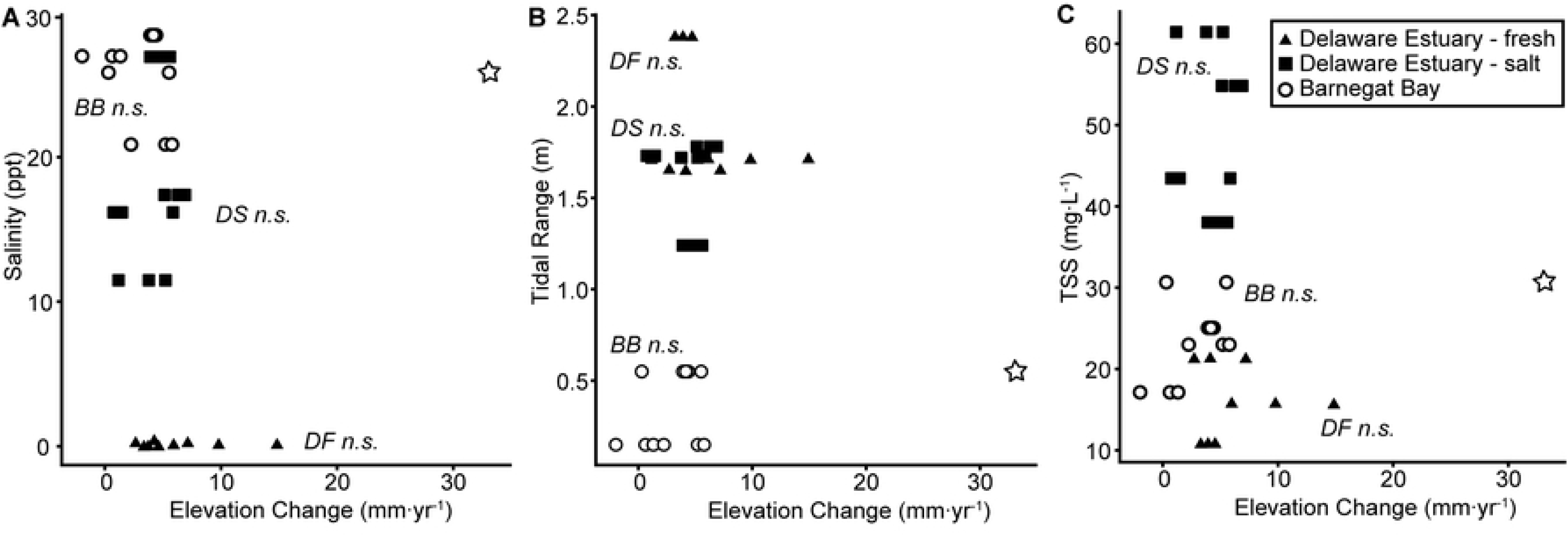
Accretion and surface elevation change for (A) the tidal freshwater marshes in the Delaware River and (B) salt marshes in the Delaware Bay. Horizontal dashed lines are LT SLR (1969–2018), ST SLR (2000–2018), or MHWR derived from the nearest NOAA tide gages (superscripts denote tidal station used for each site, north to south: (a) Philadelphia, (b) Reedy Point, (c) Cape May, and (d) Lewes. Error bars are standard error.

**Fig 5.**
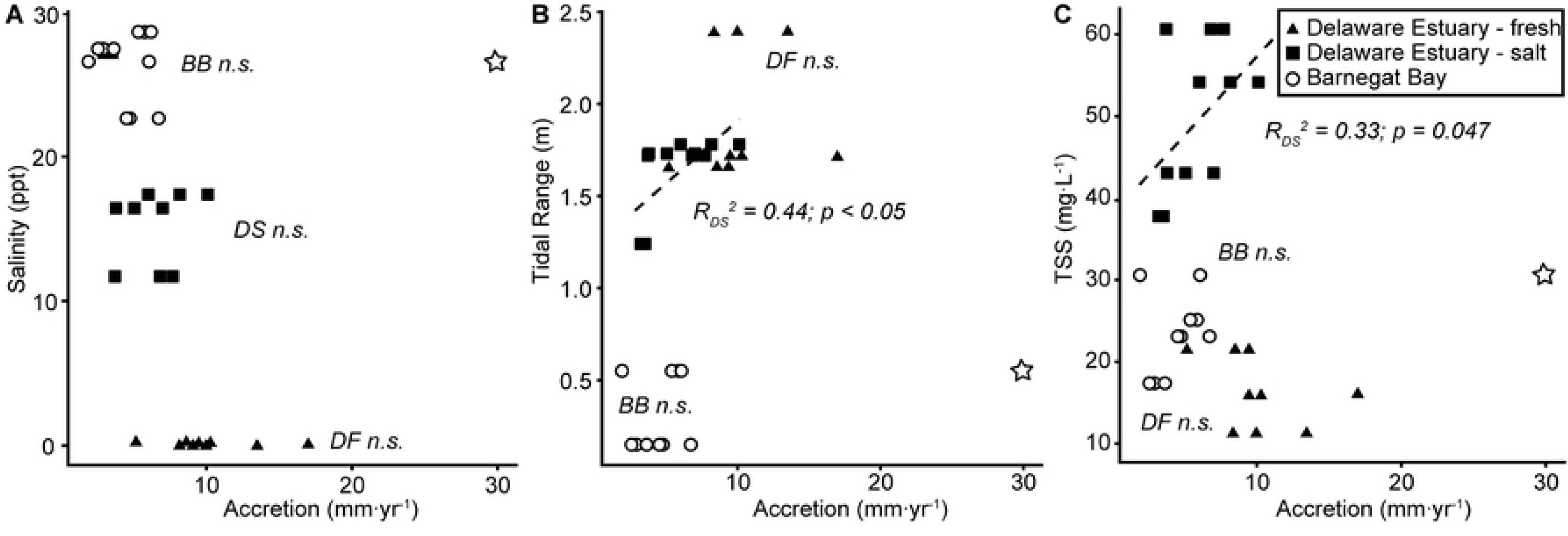
Accretion (MHs) and surface elevation change (SETs) for Barnegat Bay. Horizontal dashed lines are LT SLR (1969–2018), ST SLR (2000–2018), or MHWR derived from NOAA’s Atlantic City tide gage. Dispersal of the sediment across Dinner Point Creek SET 1 caused anomalously high annual mean accretion and elevation change rates (>25 mm yr^-1^) and so, those data were excluded. Error bars are standard error.

Associations among environmental parameters (salinity, tidal range, surface elevations, and water column suspended sediments) and accretion or elevation change rates were estuary-dependent (Figs 6–8). In the Delaware Estuary, maximum accretion rates occurred in tidal freshwater marshes, where suspended sediment concentrations were low and tidal ranges were large. While accretion rates were positively associated with tidal range (*R*^2^= 0.44, *p*<0.05) and water column suspended sediments (*R*^2^ = 0.33, *p*<0.05) in Delaware Estuary salt marshes, this association was not significant for freshwater marshes or salt marshes in Barnegat Bay. In salt marshes of the Delaware Estuary, subsurface process rates were more negative (subsidence) where salinities were lower (*R*^2^ = 0.42, *p*<0.05) and tidal ranges greater (*R*^*2*^ = 0.42, *p*<0.05).

**Fig 6.**
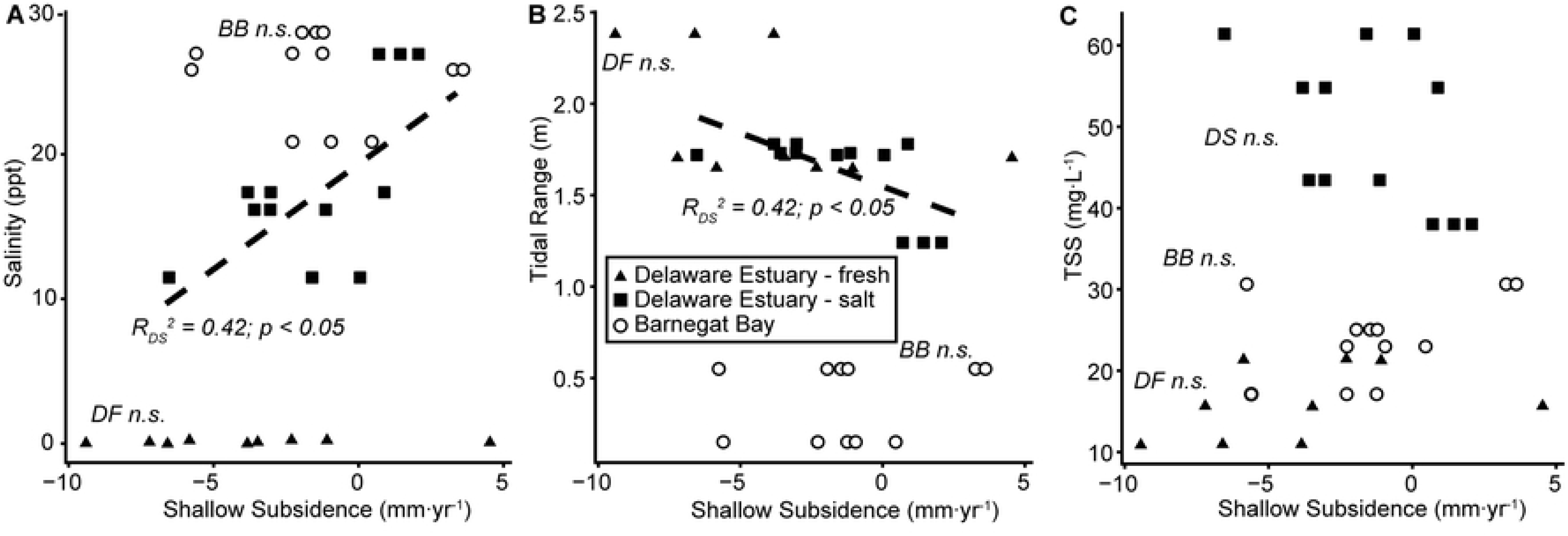
Relationship between elevation change and (A) salinity, (B) tidal range, and (C) total suspended solids (TSS). Solid triangles are freshwater tidal marshes in the Delaware River (DF) and solid squares are salt marshes in the Delaware Bay (DS). Hollow circles are salt marshes in the Barnegat Bay (BB). Dispersal of the sediment across Dinner Point Creek SET 1 as part of construction of mosquito control ponds caused anomalously high (>25 mm yr^-1^) values for accretion (hollow stars) and so, those data were excluded from analyses.

**Fig 7.**
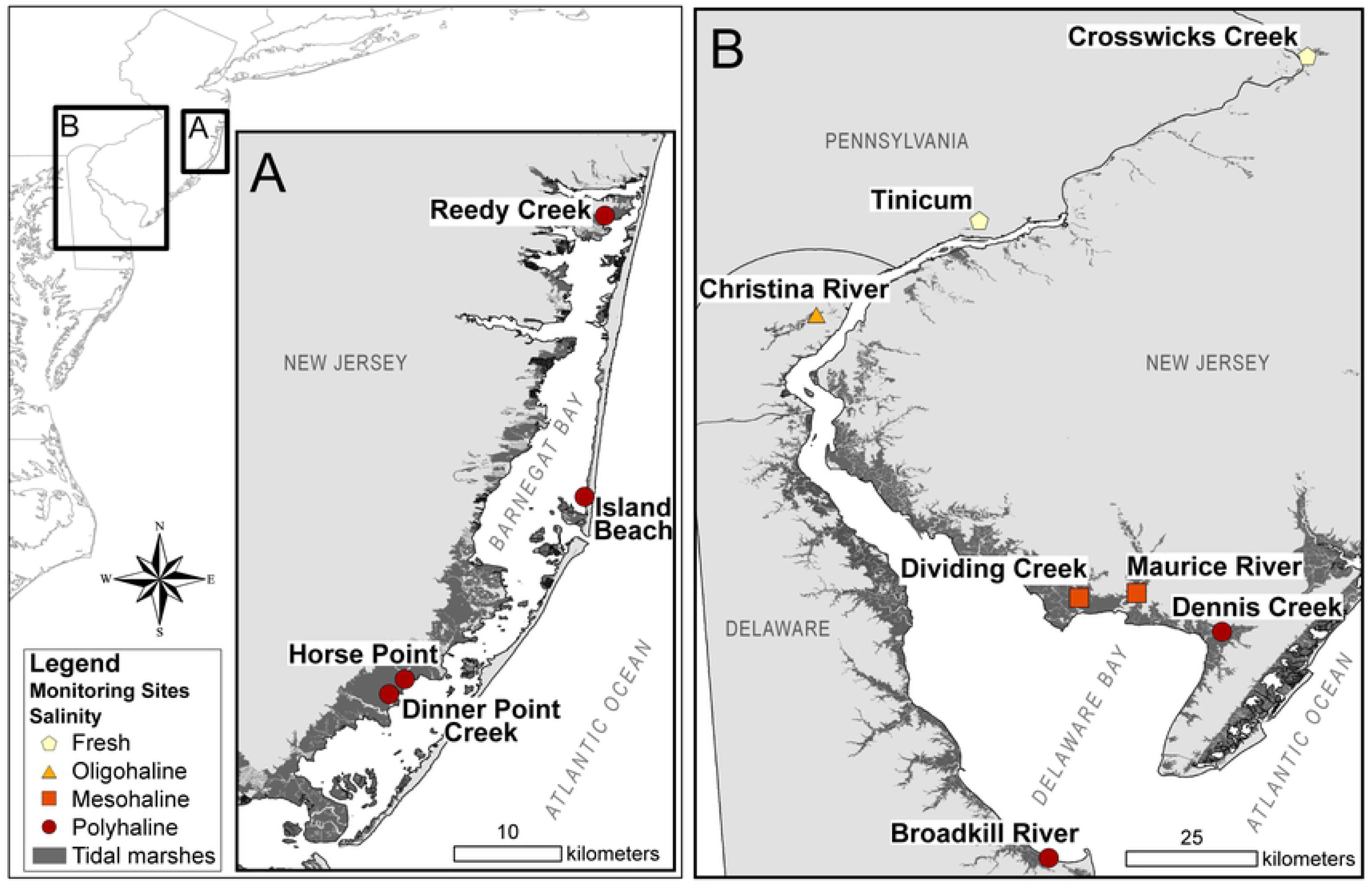
Relationship between sediment accretion and (A) salinity, (B) tidal range, and (C) total suspended solids (TSS). Solid triangles are freshwater tidal marshes in the Delaware River (DF) and solid squares are salt marshes in the Delaware Bay (DS). Hollow circles are salt marshes in the Barnegat Bay (BB). Data from Dinner Point Creek SET 1 (hollow stars) were excluded from these analyses.

**Fig 8.**
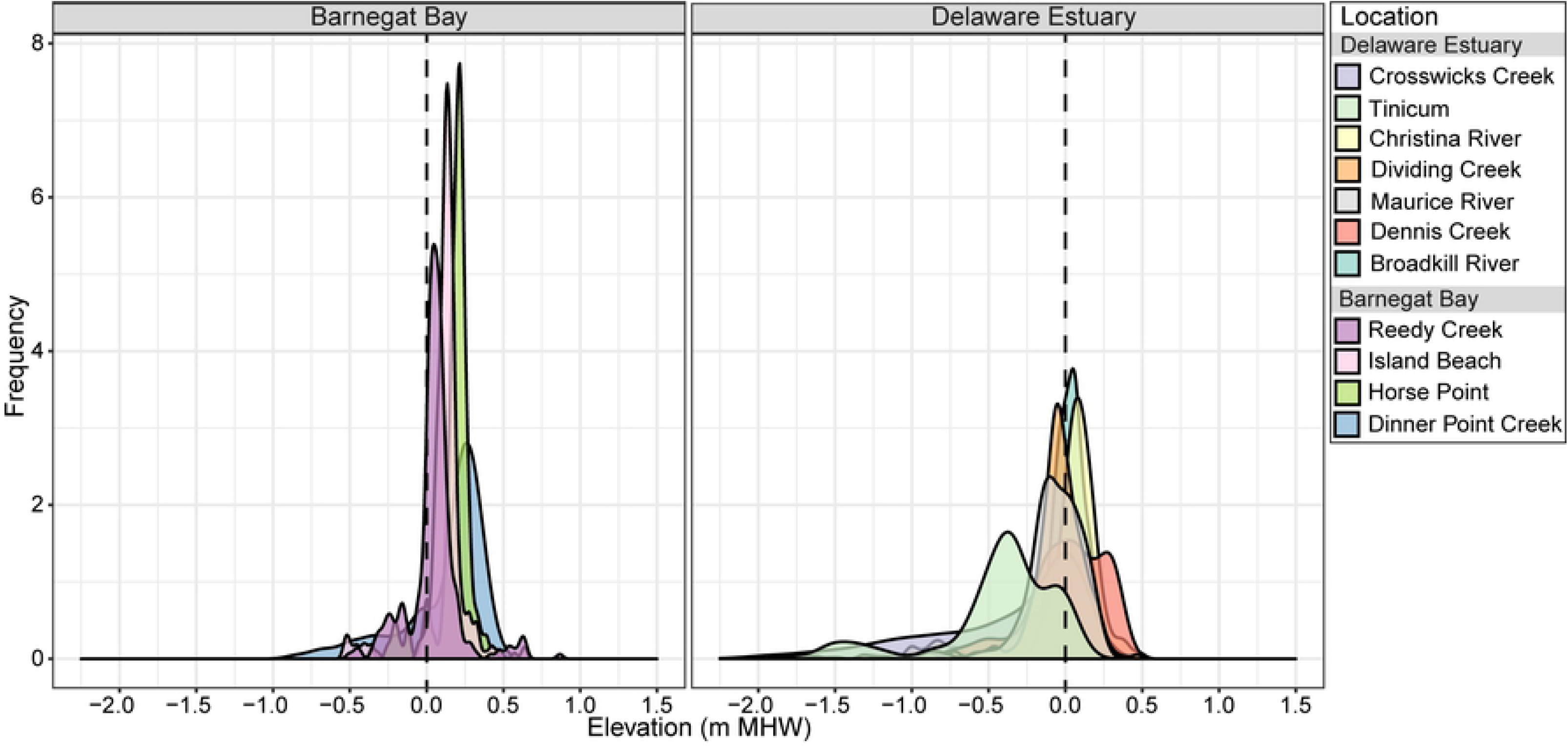
Relationship between shallow subsidence and A) salinity, (B) tidal range, and (C) total suspended solids (TSS). Solid triangles are freshwater tidal marshes in the Delaware River (DF), solid squares are salt marshes in the Delaware Bay (DS), and hollow circles are salt marshes in the Barnegat Bay (BB). Solid triangles are freshwater tidal marshes in the Delaware River, solid squares are salt marshes in the Delaware Bay, and hollow circles are salt marshes in the Barnegat Bay. R_D_^2^ is the Delaware Estuary and R_B_^2^ is for Barnegat Bay.

Sediment placement by OMWM at Dinner Point Creek SET 1 increased absolute elevations by more than 18 cm, but elevations have declined by 10 cm since 2013 (Supporting Information). Rates of elevation change were 4.9 mm yr^-1^ (*R*^*2*^= 0.93, *p* < 0.05) before placement, then declined to −13.5 mm yr^-1^ (*R*^*2*^=0.83, *p* < 0.001) after placement. These rates are not representative of natural accretion or elevation change so the Dinner Point Creek SET 1 data point was removed from elevation change and accretion analyses. Rates of shallow subsidence at this location, however, were not anomalous so we retained those values.

## Discussion

Few of the studied tidal marshes had rates of elevation change equal to or greater than LT or ST LSLR, suggesting that regional marsh loss may be associated with deficits in vertical accretion. Although accretion rates were comparable to LT LSLR, shallow subsidence offset the effects of surface accretion, and thus few sites had elevation change rates that “kept pace” with LT LSLR. From 1996–2010, tidal marsh area in Barnegat Bay and the Delaware Estuary declined by 199 ha yr^-1^ [35] and 194 ha yr^-1^ [36], respectively. Tidal marsh loss in Maryland [40], New York City [58], and Rhode Island [42] have been similarly linked to elevation changes less than LSLR. Losses, coupled with ongoing interior marsh loss and deterioration [27–28, 34, 37], indicate that many marshes in this region are unable to maintain elevations relative to accelerating sea level rise.

Rates of elevation change, accretion, and shallow subsidence varied across our study sites, which included tidal marshes in a coastal lagoon as well as in tidal freshwater and saline portions of a large coastal plain estuary. In the Delaware Estuary, rates of accretion and subsurface processes varied between salt (median values of 5.5 mm yr^-1^ for accretion and −1.4 mm yr^-1^ for shallow subsidence) and freshwater marshes (median values of 9.5 mm yr^-1^ for accretion and −3.8 mm yr^-1^ for shallow subsidence). Differences in dominant vegetation structure and sediment sourcing contribute to higher rates of accretion in tidal freshwater marshes compared to salt marshes, as others have previously discussed [59–61]. In Barnegat Bay and Delaware Estuary salt marshes, magnitudes of elevation change (median values of 4.9 and 4.1 mm yr^-1^, respectively) and accretion (median values of 5.5 and 5.1 mm yr^-1^, respectively) were comparable, despite higher salinities, lower suspended sediment concentrations, and smaller tidal ranges in Barnegat Bay. Previous findings suggest that in addition to sedimentation, organic production plays an important role in accretion for Barnegat Bay salt marshes [31].

As rates of elevation change and accretion rates are higher at lower elevations, it can be problematic to discern drowning or vulnerability to open water conversion without added context of elevation capital [58, 62]. For instance, accretion and elevation change rates at Tinicum mostly exceeded ST SLR, yet, these marshes sit low in the tidal frame (−0.47 m MHW), and they flooded 39% of the time during the 2016 growing season. As such, rising flood frequencies may drown vegetation at Tinicum before marshes that have greater elevation capital, but lower accretion rates. Similarly, in Barnegat Bay, Reedy Creek had lower elevation capital (0.05 m MHW) and flooding was frequent (67%), despite elevation change and accretion rates that were approximately equal with LT SLR. For tidal marshes with more elevation capital, like Horse Point (0.18 m MHW), lower elevation change or accretion rates likely reflect less sediment delivery due to less frequent tidal flooding. Thus, high accumulation or high elevation gains – rather than a symptom of high SLR resilience – may instead reflect submergence due to positive feedbacks between flooding frequency, sediment deposition, and high water column suspended sediment concentrations liberated by degrading or fragmenting wetlands.

Elevation processes, specifically accretion, do not wholly capture tidal marsh vulnerability to drowning or loss as other sediment dynamics also contribute to tidal marshes stability. In two Maryland-based studies, Ganju et al. [63–64] found that net sediment export likely drove deterioration and instability of tidal marshes at Blackwater River, despite abundant suspended sediment and seemingly adequate accretion rates relative to sea level rise. Thus, Ganju et al. surmised that the release of organic marsh sediments through the deterioration process might subsidize suspended sediment concentrations and sedimentation rates [64]. In a follow-up study, Ganju et al. monitored sediment dynamics at two sites in our monitoring network: deteriorated Reedy Creek, where marshes are fragmenting, and Dinner Point Creek, which was relatively stable with a larger proportion of vegetated marsh area [65]. Those results similarly found that deterioration at Reedy Creek correlated with larger net export of sediments compared to Dinner Point Creek. From our results, rates of accretion and elevation change at Reedy Creek ranged from 4.52–6.72 mm yr^-1^ and 2.24–5.77 mm yr^-1^, respectively. Accretion and elevation change rates at Horse Point, located adjacent to Dinner Point Creek yet undisturbed from pond excavation, ranged from 5.39–5.87 mm yr^-1^and 3.92–4.40 mm yr^-1^, respectively. Our findings ultimately support previous assertions that high rates of sediment accretion or elevation do not protect against marsh loss if they are a symptom of marsh degradation and fragmentation.

Although it is convention to compare tidal marsh elevation changes to LT LSLR trends to determine if accumulation debts are accruing [51], we find that ST LSLR or MHW trends more adequately represented changing inundation regimes in tidal marshes. Analysis of water levels suggest that SLR in our region was 44% greater during the 2000–2018 period than it was over the last 50 years (1969–2018), and in addition, rates of MHW rise over the past 19 years were nearly double rates of LT SLR (87% greater) (Table 3). In this study, ST LSLR and MHWR did not increase proportionately with LT LSLR across sites, so local conditions may contribute to how inundation patterns change in response to accelerating LSLR. A previous study on tidal datum changes in the U.S. similarly found that mean high water rose faster than mean sea level at tidal gages in Atlantic City, NJ, Cape May, NJ, and Lewes, DE [66]. If local MHW rises faster than LSLR, comparing LT LSLR to elevation change rates could lead to underestimations of tidal marsh SLR vulnerability. Although differences between rates of elevation change and ST SLR or MHW presented in this study were concerning, these differences could also represent lags or shifts in tidal marsh elevation dynamics [16] that were not captured in this 8-year dataset, which only further monitoring might elucidate.

Broadly, our results serve as a baseline to evaluate how patterns between elevation change and inundation become detrimental to tidal wetlands as SLR accelerates. Elevation changes relative to LSLR dynamically associate with tidal marsh elevation capital, and so, accretion is likely to increase with rising water levels. Although increased flooding facilitates greater accretion, temporal lags may exist between increasing depth and duration of floods and cumulative sediment accumulation [16]. Tidal marsh plants are most productive within a specific range of elevations relative to local tidal datum [11], so delays in elevation building by sedimentation leads to reduced plant production as rapidly rising water levels surpass optimal growth elevations. Indeed, many recent studies have attributed marsh losses to plant intolerance to excessive flooding caused by rising water levels [29, 67–70]. Subsequent vegetated area losses cause further destabilization through net sediment export [65]. A portion of our sites had accretion rates congruent with LT SLR and nearly all were well below ST SLR, thus we posit that sedimentation paces dynamically with LT SLR but lags behind ST SLR so that biological limitations associated with plant flood tolerance drives marsh deterioration or drowning. In future studies, it may be more pragmatic to ask whether marshes are “catching up” with SLR, rather than intrinsically “keeping pace.”

## Conclusions

Rates of elevation change in most (96%) tidal marshes across the Delaware Estuary and Barnegat Bay have not kept pace with recent (2000–2018) rising water levels. Accumulation deficits, relative to LT LSLR (1969–2018), varied from −4.12 to +10.8 mm yr^-1^ in the Delaware Estuary and −6.59 to +1.08 mm yr^-1^ in Barnegat Bay. Overall, rates of elevation change, sediment accumulation, and shallow subsidence varied between estuaries and across sites. Relationships among these rates and salinity, water column suspended sediments, and tidal range differed by geography. We found that more recent water level information was useful in determining inundation frequency more precisely. Due to the importance of sustained tidal marsh acreage to coastal communities across the U. S. Mid Atlantic, we hope these data provide much-needed context for future intervention efforts focused on preventing losses due to rising sea levels in our region.

## Acknowledgements

We are immensely grateful for all support provided over the years by our numerous partners across Delaware, Pennsylvania, and New Jersey. We wish to thank Kathleen Drake, Dorina Frizzera, Irene Purdy, and Kathleen Walz for their support. We also thank Melanie Mills, Erin Reilly, and Jessie Buckner for their early contributions to this work.

## Supporting Information

**S1 Table. Data collected for calculation of empirical tidal datums.**

**S2 Table. Tidal datums calculated from 2000–2018**.

**S3 Table. Tidal datums calculated for the National Tidal Datum Epoch (1983–2001)**.

**S4 Table. Tidal datums from NOAA Vdatum**.

**S5 Table. Error between NTDE datum and Vdatum (%)**.

**S6 Table. Values of accretion, surface elevation change, shallow subsidence, and comparisons to long-term (LT; 1969–2018) local sea level rise (LSLR), short-term (ST; 2000–2018) LSLR, and mean high water rise (MHWR; 2000– 2018)(see Table 4 for those rate values)**.

Standard errors are in parenthesis. To be conservative, when rates of accretion or elevation change exceeded water level rise rates (LT LSLR, ST LSLR, or MHWR) by at least one standard error, we assigned it *yes* (bold Y, boxes gray shaded) for keeping pace; otherwise, we assigned it *no* (N) for not keeping pace.

**Fig S1. Trends of elevation change at Dinner Point Creek SET 1 before and after it was abruptly buried with sediment as part of the adjacent construction of a mosquito control pond (i**.**e. OMWM)**. Between 2011 and 2013, marsh elevation increased at a rate of 4.92 mm y^-1^; 2013 to the present, elevation decreased at a rate of 13.5 mm y^-1^. Error bars are standard error.

## References

1. Chmura GL, Anisfeld SC, Cahoon DR, Lynch JC. Global carbon sequestration in tidal, saline wetland soils. Global Biogeochem Cycles 2003;17:12. https://doi.org/111110.1029/2002gb001917.

2. Loomis MJ, Craft CB. Carbon Sequestration and Nutrient (Nitrogen, Phosphorus) Accumulation in River-Dominated Tidal Marshes, Georgia, USA. Soil Sci Soc Am J 2010;74:1028. https://doi.org/10.2136/sssaj2009.0171.

3. Carr EW, Shirazi Y, Parsons GR, Hoagland P, Sommerfield CK.. Modeling the Economic Value of Blue Carbon in Delaware Estuary Wetlands: Historic Estimates and Future Projections. J Environ Manage 2018;206:40–50. https://doi.org/10.1016/j.jenvman.2017.10.018.

4. Narayan S, Beck MW, Wilson P, Thomas CJ, Guerrero A, Shepard CC, et al. The Value of Coastal Wetlands for Flood Damage Reduction in the Northeastern USA. Sci Rep 2017;7:1–12. https://doi.org/10.1038/s41598-017-09269-z.

5. Arkema KK, Guannel G, Verutes G, Wood S a., Guerry A, Ruckelshaus M, et al. Coastal habitats shield people and property from sea-level rise and storms. Nat Clim Chang 2013;3:913–8. https://doi.org/10.1038/nclimate1944.

6. Sutton-Grier AE, Wowk K, Bamford H. Future of our coasts: The potential for natural and hybrid infrastructure to enhance the resilience of our coastal communities, economies and ecosystems. Environ Sci Policy 2015;51:137–48. https://doi.org/10.1016/j.envsci.2015.04.006.

7. Lathrop RG, Cole MB, Showalter RD. Quantifying the habitat structure and spatial pattern of New Jersey (U.S.A.) salt marshes under different management regimes. Wetl Ecol Manag 2000;8:163–172.

8. Nikitina DL, Kemp AC, Horton BP, Vane CH, van de Plassche O, Engelhart SE. Storm erosion during the past 2000 years along the north shore of Delaware Bay, USA. Geomorphology 2014;208:160–72. https://doi.org/10.1016/j.geomorph.2013.11.022.

9. Weston NB. Declining Sediments and Rising Seas: An Unfortunate Convergence for Tidal Wetlands. Estuaries and Coasts 2014;37:1–23. https://doi.org/10.1007/s12237-013-9654-8.

10. Lotze HK, Lenihan HS, Bourque BJ, Bradbury RH, Cooke RG, Kay MC, et al. Depletion, Degradation, and Recovery Potential of Estuaries and Coastal Seas. Science (80-) 2006;312:1806–9.

11. Morris JT, Sundareshwar P V., Nietch CT, Kjerfve B, Cahoon DR. Responses of coastal wetlands to rising sea level. Ecology 2002;83:2869–77. https://doi.org/10.1890/0012-9658(2002)083[2869:ROCWTR]2.0.CO;2.

12. Nyman JA, Walters RJ, Delaune RD, Patrick WH, Nyman JA, Walters RJ, et al. Marsh vertical accretion via vegetative growth. Estuar Coast Shelf Sci 2006;69:370–80. https://doi.org/10.1016/j.ecss.2006.05.041.

13. Mudd SM, Howell SM, Morris JT. Impact of dynamic feedbacks between sedimentation, sea-level rise, and biomass production on near-surface marsh stratigraphy and carbon accumulation. Estuar Coast Shelf Sci 2009;82:377–89. https://doi.org/10.1016/j.ecss.2009.01.028.

14. Cahoon DR, Guntenspergen GR. Climate Change, Sea-Level Rise, and Coastal Wetlands. Natl Wetl Newsl 2010;32.

15. D’Alpaos A, Mudd SM, Carniello L. Dynamic response of marshes to perturbations in suspended sediment concentrations and rates of relative sea level rise. J Geophys Res Earth Surf 2011;116:1–13. https://doi.org/10.1029/2011JF002093.

16. Fagherazzi S, Kirwan ML, Mudd SM, Guntenspergen GR, Temmerman S, Rybczyk JM, et al. Numerical models of salt marsh evolution: Ecological, geormorphic, and climatic factors. Rev Geophys 2012;50:1–28. https://doi.org/10.1029/2011RG000359.1.

17. Mudd SM. The life and death of salt marshes in response to anthropogenic disturbance of sediment supply. Geol Soc Am 2011;39:511–2. https://doi.org/10.1130/focus052011.1.

18. Elsey-Quirk T, Adamowicz SC. Influence of Physical Manipulations on Short-Term Salt Marsh Morphodynamics: Examples from the North and Mid-Atlantic Coast, USA. Estuaries and Coasts 2015:39:423–39. https://doi.org/10.1007/s12237-015-0013-9.

19. Engelhart SE, Peltier WR, Horton BP. Holocene relative sea-level changes and glacial isostatic adjustment of the U. S. Atlantic coast 2011; 39:751–4. https://doi.org/10.1130/G31857.1.

20. Engelhart SE, Horton BP. Holocene sea level database for the Atlantic coast of the United States. Quat Sci Rev 2012;54:12–25. https://doi.org/10.1016/j.quascirev.2011.09.013

21. Ezer T, Atkinson LP, Corlett WB, Blanco JL. Gulf Stream’s induced sea level rise and variability along the U.S. Mid-Atlantic coast. J Geophys Res Ocean 2013;118:685–97. https://doi.org/10.1002/jgrc.20091.

22. Sallenger AH, Doran KS, Howd PA. Hotspot of accelerated sea-level rise on the Atlantic coast of North America. Nat Clim Chang 2012;2:884–8. https://doi.org/10.1038/nclimate1597.

23. Church JA, Clark PU, Cazenave A, Gregory JM, Jevrejeva S, Levermann A, et al. Sea level change. Clim. Chang. 2013 Phys. Sci. Basis. Contrib. Work. Gr. I to Fifth Assess. Rep. Intergov. Panel Clim. Chang. 2013. p. 1137–216. Available from: https://www.ipcc.ch/site/assets/uploads/2018/02/WG1AR5_Chapter13_FINAL.pdf

24. Callahan JA, Benjamin P. Horton, Nikitina DL, Sommerfield CK, McKenna TE, Swallow D. Recommendation of Sea-Level Rise Planning Scenarios for Delaware: Technical Report, prepared for Delaware Department of Natural Resources and Environmental Control (DNREC) Delaware Coastal Programs. 2017. Available from: https://southbethany.delaware.gov/files/2018/11/Attachment-6-to-February-2018-Mayor-Report-Technical-Report-Regarding-SLR-Planning-Scenarios.pdf.

25. Sun H, Grandstaff D, Shagam R. Land subsidence due to groundwater withdrawal: Potential damage of subsidence and sea level rise in southern New Jersey, USA. Environ Geol 1999;37:290–6. https://doi.org/10.1007/s002540050386.

26. Smith JAM, Hafner SF, Niles LJ. The Impact of Past Management Practices on Tidal Marsh Resilience to Sea Level Rise in the Delaware Estuary. Ocean Coast Manag 2017;149:33–41. https://doi.org/10.1016/j.ocecoaman.2017.09.010.

27. Kearney MS, Rogers AS, Townshend JRG, Rizzo E, Stutzer D, Stevenson JC, et al. Landsat imagery shows decline of coastal marshes in Chesapeake and Delaware Bays. Eos, Trans Am Geophys Union 2002;83:173. https://doi.org/10.1029/2002EO000112.

28. Hartig EK, Gornitz V, Kolker A, Mushacke F, Fallon D. Anthropogenic and Climate-Change Impacts on Salt Marshes of Jamaica Bay, New York City. Wetlands 2002;22:71–89. https://doi.org/10.1672/0277-5212(2002)022[0071:AACCIO]2.0.CO;2.

29. Watson EB, Raposa KB, Carey JC, Wigand C, Warren RS. Anthropocene Survival of Southern New England’s Salt Marshes. Estuaries and Coasts 2016. https://doi.org/10.1007/s12237-016-0166-1.

30. Elsey-Quirk T. Impact of Hurricane Sandy on salt marshes of New jersey. Estuar Coast Shelf Sci 2016;183:235–48. https://doi.org/10.1016/j.ecss.2016.09.006.

31. Elsey-Quirk T, Unger V. Geomorphic influences on the contribution of vegetation to soil C accumulation and accretion in *Spartina alterniflora* marshes. Biogeosciences 2018;15:379–97. https://doi.org/10.5194/bg-15-379-2018.

32. Ferrigno F, Widjeskog L, Toth S. Pittman-Robertson Report Proj. W-53-R-1, Job I-G. In: Marsh Destruction, New Jersey Department of Environmental Protection. 1973.

33. Field RT, Philipp KR. Vegetation changes in the freshwater tidal marsh of the Delaware Estuary. Wetl Ecol Manag 2000;8: 79–88. https://doi.org/10.1023/A:1008480116062.

34. Kearney MS, Riter JCA. Inter-annual variability in Delaware Bay brackish marsh vegetation, USA. Wetl Ecol Manag 2011;19:373–88. https://doi.org/10.1007/s11273-011-9222-6.

35. Haaf L, Kreeger D, Homsey A. Chapter 5.2 - Intertidal marsh In: Technical Report for the Delaware Estuary and Basin. Partnership for the Delaware Estuary. PDE Report No. 17-07. 2017: 177–193. Available from: http://www.delawareestuary.org/data-and-reports/state-of-the-estuary-report/.

36. Buckner J, Haaf L, Padeletti A, Kreeger D, Maxwell-Doyle M. Trends in Salt Marsh Acreage for Barnegat Bay. Barnegat Bay Partnership, Toms River, NJ. 2018:1–16.

37. Erwin RM, Cahoon DR, Prosser DJ, Sanders GM, Hensel P. Surface elevation dynamics in vegetated *Spartina* marshes versus unvegetated tidal ponds along the Mid-Atlantic coast, USA, with implications to waterbirds. Estuaries and Coasts 2006;29:96–106. https://doi.org/10.1007/BF02784702.

38. Delgado P, Hensel PF, Swarth CW, Ceroni M, Boumans R. Sustainability of a Tidal Freshwater Marsh Exposed to a Long-term Hydrologic Barrier and Sea Level Rise: A Short-term and Decadal Analysis of Elevation Change Dynamics. Estuaries and Coasts 2013;36:585–94. https://doi.org/10.1007/s12237-013-9587-2.

39. Cadol D, Engelhardt K, Elmore A, Sanders G. Elevation-dependent surface elevation gain in a tidal freshwater marsh and implications for marsh persistence. Limnol Oceanogr 2014;59:1065–80. https://doi.org/10.4319/lo.2014.59.3.1065.

40. Beckett LH, Baldwin AH, Kearney MS. Tidal marshes across a chesapeake bay subestuary are not keeping up with sea-level rise. PLoS One 2016;11:1–12. https://doi.org/10.1371/journal.pone.0159753.

41. Carey JC, Raposa KB, Wigand C, Warren RS. Contrasting decadel-scale changes elevation and vegetation in in two Long Island Sound salt marshes. Estuaries and Coasts 2017;40:651–61. https://doi.org/10.1007/s12237-015-0059-8.Contrasting.

42. Raposa KB, Cole ML, Burdick DM, Ernst T, Adamowicz SC. Elevation change and the vulnerability of Rhode Island (USA) salt marshes to sea-level rise. Reg Environ Chang 2017;71:398–97. https://doi.org/10.1007/s10113-016-1020-5.

43. Pennsylvania Natural Heritage Program. A Natural Heritage Inventory of Philadelphia County, Pennsylvania. 2008:1–248. Available from: http://www.naturalheritage.state.pa.us/cnai_pdfs/philadelphia_county_nhi_2008_web.pdf

44. Ferren WRJ, Schuyler AE. Intertidal Vascular Plants of River Systems near Philadelphia. Proc Acad Nat Sci Philadelphia 1980;132:86–120.

45. Defne Z, Ganju NK. Quantifying the Residence Time and Flushing Characteristics of a Shallow, Back-Barrier Estuary: Application of Hydrodynamic and Particle Tracking Models. Estuaries and Coasts 2014;38:1719–34. https://doi.org/10.1007/s12237-014-9885-3.

46. Barnegat Bay Partnership. State of the Bay. 2016:1–80. Available from: https://www.barnegatbaypartnership.org/wp-content/uploads/2017/08/BBP_State-of-the-Bay-book-2016_forWeb-1.pdf.

47. Powell, E. P. The Effect of Open Marsh Water Management Practices on the Carbon Balance of Tidal Marshes in Barnegat Bay, New Jersey. Master Thesis. Drexel University. 2018. Available from: http://hdl.handle.net/1860/idea:7834

48. Cahoon DR, Lynch JC, Perez BC, Segura B, Holland RD, Stelly C, et al. High-Precision Measurements of Wetland Sediment Elevation: II. The Rod Surface Elevation Table. J Sediment Res 2002;72:734–9. https://doi.org/10.1306/020702720734.

49. Lynch JC, Hansel P, Cahoon DR. The Surface Elevation Table and Marker Horizon Technique A Protocol for Monitoring Wetland Elevation Dynamics. Natural Resource Report NPS/NCBN/NRR—2015/1078. 2015. Available from: https://irma.nps.gov/DataStore/Reference/Profile/2225005.

50. Cahoon DR, Turner RE. Accretion and canal impacts in a rapidly subsiding wetland II. Feldspar marker horizon technique. Estuaries 1989;12:260–8.

51. Cahoon DR. Estimating Relative Sea-Level Rise and Submergence Potential at a Coastal Wetland. Estuaries and Coasts 2015;38:1077–84. https://doi.org/10.1007/s12237-014-9872-8.

52. Henning, W. User Guidelines for Single Base Real Time GNSS Positioning. National Geodetic Survey: Silver Spring, MD, USA, 2011:1–138.

53. National Oceanic and Atmospheric Administration. Computational Techniques for Tidal Datums Handbook. 2003. Available from: https://tidesandcurrents.noaa.gov/publications/Computational_Techniques_for_Tidal_Datums_handbook.pdf

54. Swanson RL. Variability of tidal datums and accuracy in determining datums from short series of observations. National Oceanic and Atmospheric Administration Tech. Rep. NOS 64. 1974:1–41.

55. Wickham H. ggplot2: Elegant Graphics for Data Analysis. Springer-Verlag New York. 2009.

56. R Core Team. R: A language and environment for statistical computing. R Foundation for Statistical Computing, Vienna, Austria. 2019. Available from: http://www.R-project.org/

57. National Oceanic and Atmospheric Administration Tides & Currents. Station IDs 8545240, 8551910, 8536110, and 8557380. Available from https://tidesandcurrents.noaa.gov/sltrends/

58. Cahoon DR, Lynch JC, Roman CT, Schmit JP, Skidds DE. Evaluating the Relationship Among Wetland Vertical Development, Elevation Capital, Sea-Level Rise, and Tidal Marsh Sustainability. Estuaries and Coasts 2019;42:1–15. https://doi.org/10.1007/s12237-018-0448-x.

59. Ensign SH, Hupp CR, Noe GB, Krauss KW, Stagg CL. Sediment Accretion in Tidal Freshwater Forests and Oligohaline Marshes of the Waccamaw and Savannah Rivers, USA. Estuaries and Coasts 2014;37:1107–19. https://doi.org/10.1007/s12237-013-9744-7.

60. Ensign SH, Noe GB, Hupp CR, Skalak KJ. Head-of-tide bottleneck of particulate material transport from watershed to estuaries. Geophys Res Lett 2015;42:671–9. https://doi.org/10.1002/2015GL066830.Received.

61. Palinkas CM, Engelhardt KAM. Influence of Inundation and Suspended-Sediment Concentrations on Spatiotemporal Sedimentation Patterns in a Tidal Freshwater Marsh. Wetlands 2019;39:507–20. https://doi.org/10.1007/s13157-018-1097-3.

62. Cahoon DR, Reed DJ, Day JW. Estimating shallow subsidence in microtidal salt marshes of the southeastern United States: Kaye and Barghoorn revisited. Mar Geol 1995;128:1–9. https://doi.org/10.1016/0025-3227(95)00087-F.

63. Ganju NK, Nidzieko NJ, Kirwan ML. Inferring tidal wetland stability from channel sediment fluxes: Observations and a conceptual model. J Geophys Res Earth Surf 2013;118:2045–58. https://doi.org/10.1002/jgrf.20143.

64. Ganju NK, Kirwan ML, Dickhudt PJ, Guntenspergen GR, Cahoon DR, Kroeger KD. Sediment transport-based metrics of wetland stability. Geophys Res Lett 2015;42:7992–8000. https://doi.org/10.1002/2015GL065980.

65. Ganju NK, Defne Z, Kirwan ML, Fagherazzi S, D’Alpaos A, Carniello L, et al. Spatially integrative metrics reveal hidden vulnerability of microtidal salt marshes. Nat Commun 2017;8:14156. https://doi.org/10.1038/ncomms14156.

66. Flick RE, Murray JF, Ewing LC. Trends in United States Tidal Datum Statistics and Tide Range: A Data Report Atlas. Scripps Institution of Oceanography (SIO) Reference Series 99-20. 1999:1–213.

67. Kirwan ML, Guntenspergen GR. Influence of tidal range on the stability of coastal marshland. J Geophys Res Earth Surf 2010;115. https://doi.org/10.1029/2009jf001400.

68. Snedden GA, Cretini K, Patton B. Inundation and salinity impacts to above- and belowground productivity in Spartina patens and Spartina alterniflora in the Mississippi River deltaic plain: Implications for using river diversions as restoration tools. Ecol Eng 2014;81:133–9. https://doi.org/10.1016/j.ecoleng.2015.04.035.

69. Watson EB, Wigand C, Davey EW, Andrews HM, Bishop J, Raposa KB. Wetland Loss Patterns and Inundation-Productivity Relationships Prognosticate Widespread Salt Marsh Loss for Southern New England. Estuaries and Coasts 2016;40:662–681. https://doi.org/10.1007/s12237-016-0069-1.

70. Payne AR, Burdick DM, Moore GE. Potential Effects of Sea-Level Rise on Salt Marsh Elevation Dynamics in a New Hampshire Estuary. Estuaries and Coasts 2019:1405–18. https://doi.org/10.1007/s12237-019-00589-z.

